# Independent host- and bacterium-based determinants protect a model symbiosis from phage predation

**DOI:** 10.1101/2021.07.09.451802

**Authors:** Jonathan B. Lynch, Brittany D. Bennett, Bryan D. Merrill, Edward G. Ruby, Andrew J. Hryckowian

## Abstract

Bacteriophages (phages) are diverse and abundant constituents of microbial communities worldwide, and are capable of modulating bacterial populations in diverse ways. Here we describe a novel phage, ϕHNL01, which infects the marine bacterium *Vibrio fischeri*. We use culture-based approaches to demonstrate that mutations in the exopolysaccharide locus of *V. fischeri* render this bacterium resistant to infection by ϕHNL01, highlighting the extracellular matrix as a key determinant of phage tropism in this interaction. Additionally, using the natural symbiosis between *V. fischeri* and the squid *Euprymna scolopes*, we show that during colonization, *V. fischeri* is protected from phage present in the ambient seawater. Taken together, these findings shed light on independent yet synergistic host- and bacterium-based strategies for resisting symbiosis-disrupting phage predation, and present important implications for understanding these strategies in the context of host-associated microbial ecosystems.

## Introduction

Bacteriophages (phages) are viruses that infect bacteria, influencing broad areas of bacterial physiology, ecology, and interspecies relationships. These effects can manifest through bacterial-host cell lysis, horizontal gene transfer, or bacterial resistance mechanisms to viral infection, and they can have positive, negative, or neutral effects on bacterial fitness (Brum et al., 2015; Matilla et al., 2014; Roossinck, 2011; Suttle, 2007). Phages are abundant across diverse environments, including the mammalian gastrointestinal tract, rhizosphere, and oceanic plankton, where they can dramatically alter not only bacterial physiology but also wider environmental nutrient cycling (Al-Shayeb et al., 2020; Breitbart et al., 2018; Dion et al., 2020; Reyes et al., 2010).

The potential to impact broader community structuring has ignited great interest in the role that phages could play in shaping eukaryote-associated microbial communities known as the microbiota. In these associations, phages could select for or against certain bacterial taxa, alter activity of bacteria-derived infection factors, or influence spatial organization of the microbiota (Barr et al., 2013), enticingly suggesting “phage therapy” as a potential intervention to selectively alter the composition, and thus functions, of the microbiota.

Many details of phage biology have been revealed through the study of well-described model systems, such as *Escherichia coli*–T4, *Escherichia coli*–T7, and *Escherichia coli*–λ (Kutter et al., 2018), and *Vibrio cholerae*–ICP1, *Vibrio cholerae*–ICP2, and *Vibrio cholerae*–ICP3 phages (Yen and Camilli, 2017), among others (Ofir and Sorek, 2018). Despite these efforts, there is a dearth of knowledge about how most phages interact with their bacterial hosts in their natural environments. This knowledge gap is due both to the high diversity of phages and their target bacteria, and to the difficulty of inferring these relationships from sequence homology, bacterial-host relatedness, or relative abundances of bacterial host and phage (Dion et al., 2020; Reyes et al., 2010). For example, the breadth of the host range varies amongst phages, spanning from specific strains to entire taxonomic orders (Beumer and Robinson, 2005; Flores et al., 2011; de Jonge et al., 2019). In addition, bacterially-encoded phage-resistance strategies can be either broadly or narrowly protective (*e.g*., alteration of extracellular structures or CRISPR spacer acquisition, respectively); conversely, phages have evolved myriad ways to counter these defenses (Black, 1988; Fan et al., 2018; Morona and Henning, 1984; Parent et al., 2014; Porter et al., 2020; Wang et al., 2019; Xu et al., 2014; Zborowsky and Lindell, 2019). Bacterial evasion strategies can be constitutive or differentially expressed under particular environmental conditions (Reyes-Robles et al., 2018), further complicating an understanding of phage-bacterium interactions and highlighting the need for diverse and tractable model systems to study these associations.

In animal-bacterial symbioses, the animal host also plays a key, but understudied, role in phage dynamics. For example, phages can adhere to mucus found at epithelium-adjacent surfaces, altering phage dynamics by creating regions of high and low phage abundance and preferentially creating areas of higher phage activity (Barr et al., 2013, 2015). The animal environment can also deploy antimicrobials like immunoglobin A (IgA) that can broadly bind to and inactivate phages (Bunker et al., 2017). Indirectly, the host-tissue environment can induce differential expression of phage receptors in bacterial symbionts, changing their viral susceptibility by shifting their physiological profile (Porter et al., 2020).

While many animal microbiotas are highly diverse communities, rendering them difficult to study, some animals develop mutualisms with a narrowly defined set of bacteria that more readily enable mechanistic probing of these relationships. One well-studied example is the bioluminescence-based mutualism between the Hawaiian bobtail squid, *Euprymna scolopes*, and the marine γ-Proteobacterium *Vibrio (Aliivibrio) fischeri*. Within hours after the squid hatch from their eggs, *V. fischeri* cells from the surrounding seawater colonize a specialized symbiotic tissue known as the light organ (LO), where the bacteria multiply and reside for the approximately 9-month duration of the squid’s life (McFall-Ngai, 2014). In the lab, the colonization process is marked by an extreme bottlenecking, in which ~10^3^–10^6^ cells/mL of *V. fischeri* in the squid’s seawater are whittled down to only a few cells that initiate the colonization (Wollenberg and Ruby, 2009). These successfully colonizing bacteria are further sorted into the six distinct crypts of the LO, so that each crypt ordinarily contains one or two clonal populations of *V. fischeri* cells (Bongrand and Ruby, 2019; Speare et al., 2018; Sun et al., 2016). To ensure successful colonization of symbiosis-competent *V. fischeri* while simultaneously restricting LO access of non-*V. fischeri* microbes, *E. scolopes* employs several strategies, such as cilia-derived flow patterns, that physically control access to the LO (Nawroth et al., 2017).

Due to its tractable partnership, the squid-*Vibrio* mutualism presents an ideal system in which to study the role of phages in beneficial host-microbe associations. We isolated and characterized a novel phage with marked specificity for a symbiosis-competent *V. fischeri* strain, as well as spontaneous phage-resistant mutants of this strain. We hypothesized that the near-clonality of *V. fischeri* in the LO would predispose decimation of these symbionts upon the introduction of a virulent phage, potentially reducing or removing the resident *V. fischeri* population and undoing the previously stable host-symbiont relationship. Surprisingly, our data demonstrate that while phages prey on populations of *V. fischeri* in the ambient seawater, *V. fischeri* cells present in the LO are protected from phage predation. We posit that a host-mediated protection of symbionts from phages would provide a fitness advantage by discouraging a costly phage sweep of the symbiont population (*e.g.*, no light production in the case of the squid–*Vibrio* symbiosis) or where general microbiota stability, rather than persistent phage-driven oscillations in community structure, promotes host homeostasis. This work identifies how animal hosts and their symbiotic bacteria can respond to and protect highly specific mutualisms in the face of phage predation, and establishes a framework with which to understand the mechanisms that govern these relationships in diverse host-associated microbial ecosystems.

## Results

### Isolation and genomic characterization of ϕHNL01, and comparative analysis with *Vibrio cholerae* phage ICP1

Using a previously reported protocol (Hryckowian et al., 2020), we isolated phage ϕHNL01 from coastal Hawaiian seawater (**Figure 1A**). ϕHNL01 was propagated on *V. fischeri* ES114, the well-studied LO isolate of this species (**Figure 1B**). We assayed the bacterial-host range of ϕHNL01 by performing plaque assays on 24 additional strains of *V. fischeri*, as well as five previously described mutants of ES114 (**Table S1**). As isolates from the ES114 background were the only strains that formed visible plaques, we conclude that ϕHNL01 has a restricted base host range and that several common extracellular macromolecules do not impact its ability to infect. For example, mutation of neither the O-antigen of lipopolysaccharide, the well-studied symbiosis polysaccharide, nor the dominant outer membrane protein, OmpU, affected the ability of ϕHNL01 to infect ES114; similarly, the central trait of the symbiosis, bioluminescence, was not required for phage infection (**Table S1**). These results suggest that the specificity of ϕHNL01 for ES114 is driven at least in part by other cellular structures.

**Figure 1.**
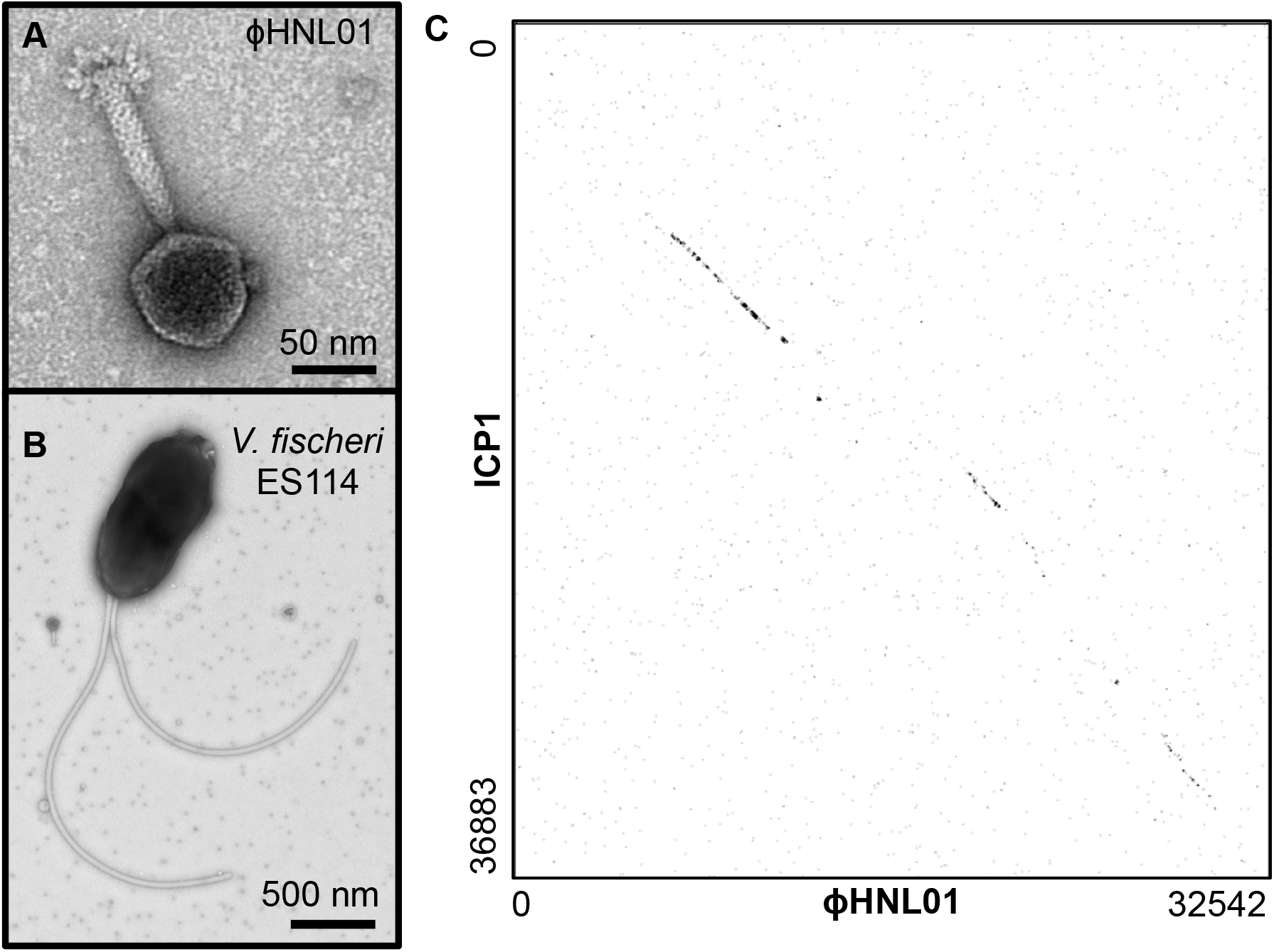
Isolation and genomic characterization of ϕHNL01. (A and B) Transmission electron micrographs of (A) ϕHNL01 and (B) *V. fischeri* ES114 and ϕHNL01. C) The amino acid sequences of all annotated protein coding ORFs in ϕHNL01 were concatenated and compared to the concatenated protein-coding ORFs in *V. cholerae* phage ICP1 with a dotplot. Numbers denote position within the concatenated amino acid sequence for each phage.

Transmission electron microscopy of ϕHNL01 demonstrated that this phage is a myovirus (e.g., tailed phage with a contractile tail; **Figures 1A, 1B**), and whole genome sequencing of ϕHNL01 revealed a 111,792 bp genome consisting of 38% GC content and 166 predicted protein-coding genes, 33 of which have conserved domains (**Figure S1, Table S2**). The virion morphology and genome size of ϕHNL01 resemble the well-studied *Vibrio cholerae*-infecting phage ICP1, a myovirus with a 125,956 bp genome (Seed et al., 2011). A relationship between the ϕHNL01 and ICP1 genomes was not evident at the nucleotide level. However, sequence similarity at the amino acid level was apparent between these two phages (**Figure 1C**), with multiple conserved proteins involved in nucleotide metabolism and virion structure (**Table S2**, **Figure S1**). Together, this resemblance suggests a more distant evolutionary history for ϕHNL01, as has been observed for phages that infect gut-resident Bacteroidetes (Guerin et al., 2018; Hryckowian et al., 2020).

### Phage predation does not impact *V. fischeri* colonization of the *E. scolopes* LO

Because phages are ubiquitous in nature and are increasingly recognized as shaping host-associated ecosystems (Koskella and Brockhurst, 2014), we sought to understand how ϕHNL01 impacts the establishment and maintenance of LO colonization, using controlled experimental approaches. First, newly hatched, uncolonized juvenile squid were colonized with *V. fischeri* ES114 for 24 hours (**Figure 2A, left**), then phages were added to the ambient seawater to a final concentration of 10^7^ plaque-forming units per milliliter (PFU mL^−1^). After 24 hours of phage exposure (*i.e*., 48 hours after initiation of bacterial colonization), the number of viable *V. fischeri* in the LO were quantified. The results highlighted that phage predation from the ambient environment does not reduce the maintenance of the normal population level of symbionts (**Figure 2A, right**; no phage = 8.25×10^4^ colony-forming units per LO (CFU/LO); plus phage = 1.55×10^5^ CFU/LO; p = 0.07). To mimic conditions that might be encountered in coastal seawater, we simultaneously exposed juvenile squid to both excess phage (at ~10^7^ PFU/mL) and *V. fischeri* cells (at ~8×10^4^ CFU/mL) and allowed colonization to proceed for 24 hours. We similarly observed no difference in colonization between the two colonization groups (**Figure 2B;** no phage = 2.1×10^5^ CFU/LO; plus phage = 1.75×10^5^CFU/mL, p = 0.78). Subsequently, homogenates from either ten *V. fischeri-*colonized squid or ten phage-exposed, *V. fischeri*-colonized squid from the colonizations represented in **Figure 2B** were tested for phage sensitivity in a soft agar overlay. ϕHNL01 still similarly infected ES114 extracted from squid, regardless of phage exposure during initiation of colonization (**Figure 2C**). Together, these data indicate that bacteria that have colonized the LO are protected from ambient phages, but that these LO-derived bacteria do not acquire permanent resistance to phages.

**Figure 2.**
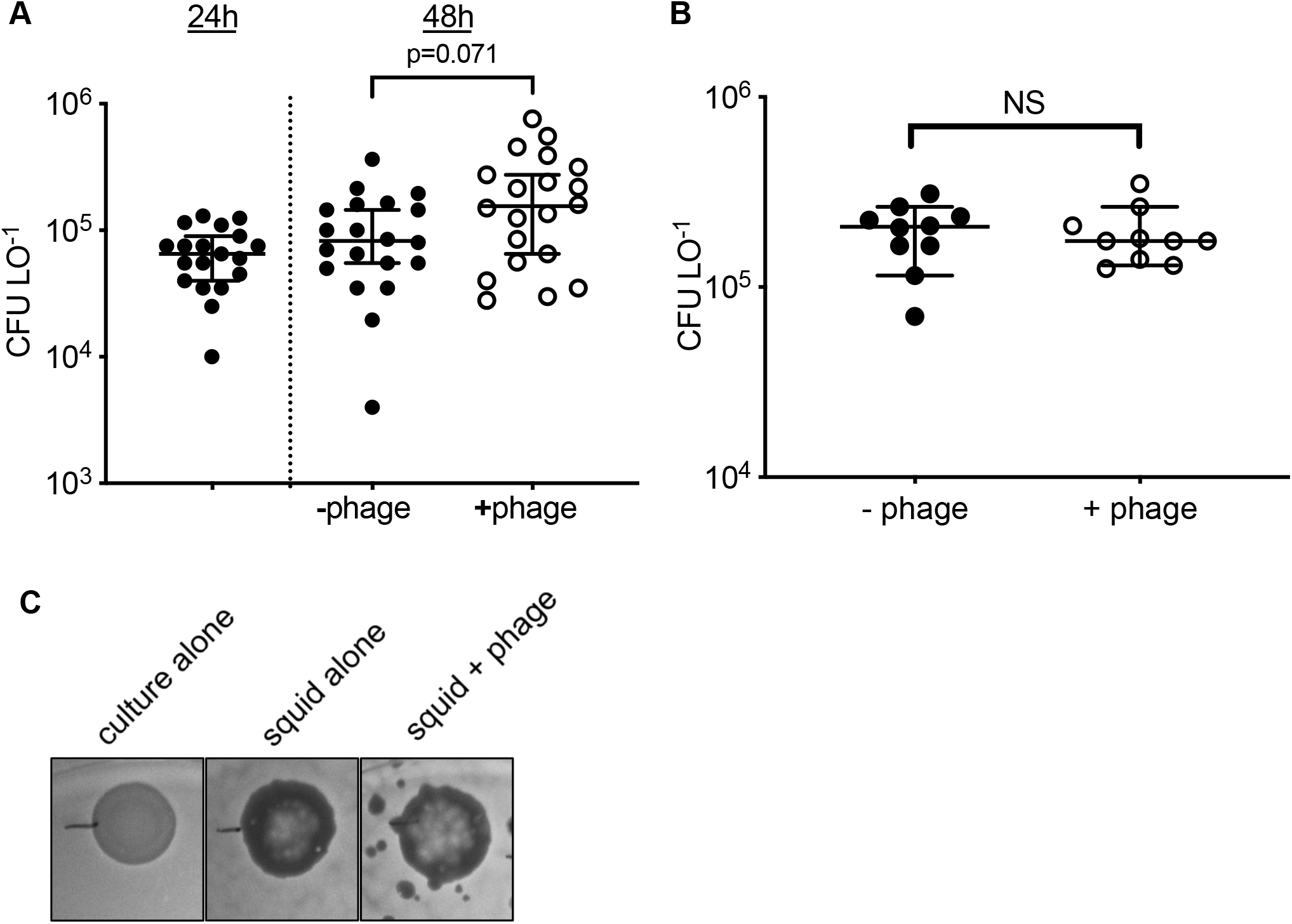
Phage predation does not reduce squid colonization, despite symbiont maintenance of viral sensitivity. A) Colony-forming units (CFU) of *V. fischeri* ES114 from squid light organ (LO) homogenates. Squid were colonized for 24 h (left) in the absence of phage, then exposed to phage, and CFU were measured at 48 h. n = 19-20 for each experimental condition. B) Squid were exposed to ϕHNL01 and *V. fischeri* at the same time and CFU LO^−1^ was measured at 24 h. n = 10 for each condition. C) ϕHNL01 spotted on top agar overlay of *V. fischeri* from an overnight culture (left), from *V. fischeri* extracted from colonized light organs (middle), or from light organs of squid concurrently exposed to *V. fischeri* and ϕHNL01 (right). Each point represents CFU from the LO of one squid. Bars represent median +/− 95% confidence interval (CI).

### Phage resistance mutations in ES114 do not impact LO colonization

In addition to the phage protection provided to *V. fischeri* by the LO (**Figure 2**), we asked whether phage resistance could be acquired *in vitro* to protect *V. fischeri* outside the squid. To address this question, we added ϕHNL01 at a range of multiplicities of infection to cultures of *V. fischeri*, and monitored changes in OD_600_ over time for 16 hours. Independent of the inoculum size or multiplicity of infection, a decrease in OD_600_ was observed within the first ~1 hour of phage exposure in cultures where phage was added, followed by ~5 hours during which bacteria were undetectable, and a subsequent outgrowth of presumably resistant bacteria (**Figure 3A**). We collected bacteria from these cultures, re-challenged them with ϕHNL01, and compared their phage sensitivity to that of cells from the phage-free cultures. As expected, bacteria that were not previously exposed to ϕHNL01 were susceptible, while in contrast, derivatives that had survived exposure to ϕHNL01 were non-susceptible, suggesting selection for heritable phage-resistance mutations (**Figure 3B**).

**Figure 3:**
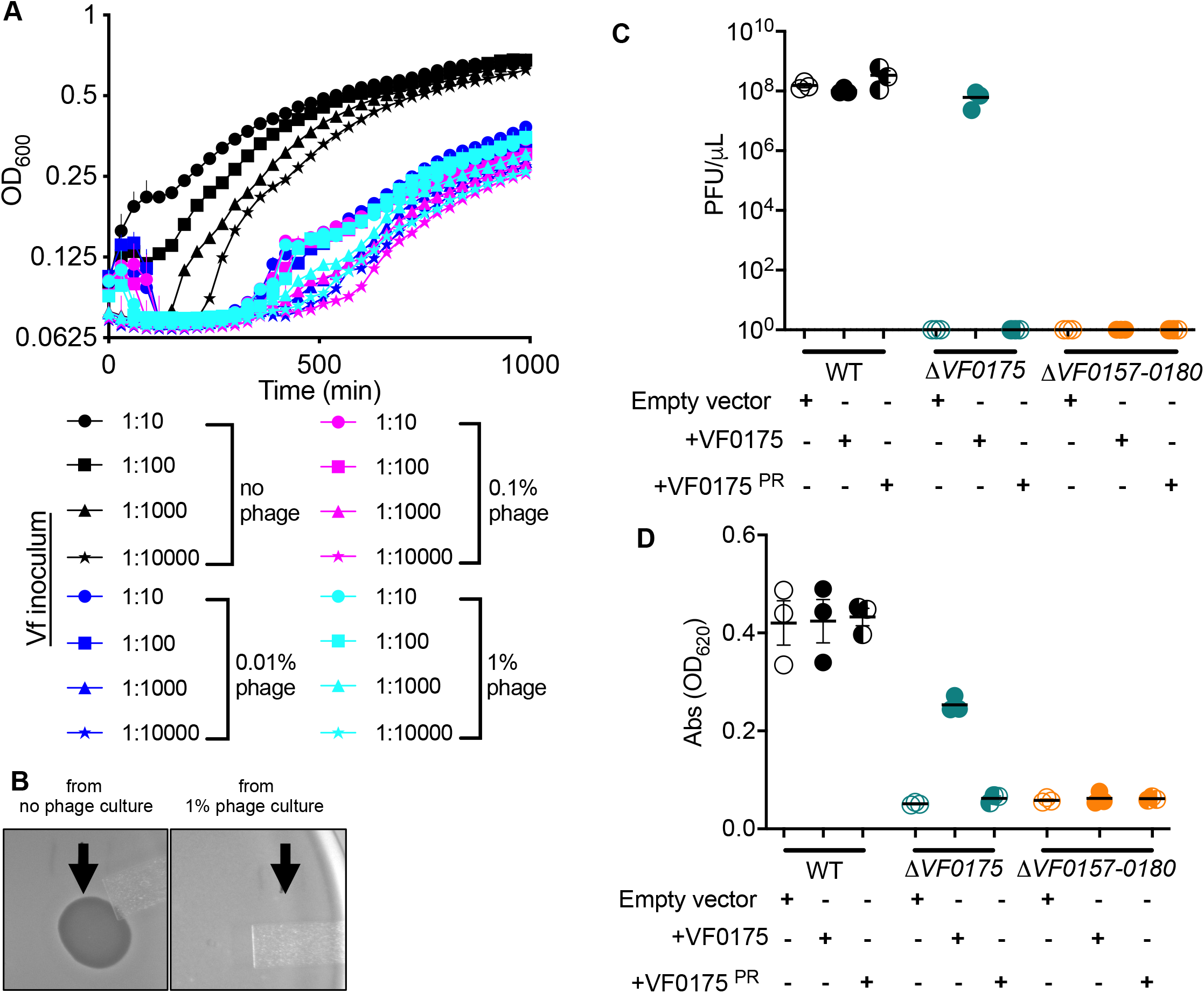
*V. fischeri* rapidly displays phage resistance during exposure to phage during *in vitro* growth. A) Growth curve of *V. fischeri* ES114 with varying dilutions of phage (0–1% by volume of ~10^11^ PFU mL^−1^ stock) and initial *V. fischeri* inoculum (1:10–1:10000 of an overnight culture). Points with error bars represent mean +/− standard error of the mean (SEM). (B) Phage spotting using stationary phase *V. fischeri* cells from cultures grown without (left) or with (right) phage. Arrows indicate location of phage spots. (C) Plaque assay measurements of ϕHNL01 infection of wild-type *V. fischeri* ES114 (black), ΔVF0175 (blue), and ΔVF0157-80 (orange) carrying empty vector (open symbols), vector with wild-type VF0175 (closed symbols), or vector with VF0175^PR^ (half-filled symbols). Points along the x-axis represent results falling below the limit of detection (1 PFU mL^−1^). (D) Alcian blue was used to stain EPS in supernatants of 24-h liquid cultures grown in minimal salts medium supplemented with 6.5 mM *N*-acetylneuraminic acid 0.05% casamino acids for wild-type *V. fischeri* ES114, ΔVF0175, and ΔVF0157-80 carrying empty vector, vector with wild-type VF0175, or vector with VF0175^PR^ (same labeling scheme as 3C). Points represent biological replicates, bars represent mean +/− SEM.

From *V. fischeri* ES114 colonies appearing in the midst of an ϕHNL01 plaque, we retrieved four individual colonies. After restreaking them to isolation, we confirmed that they were resistant to ϕHNL01 lysis in plaque assay (similar to **Figure 3B)**. We performed PCR on these isolates with ϕHNL01-specific primers to test for lysogeny, and did not observe any of the tested phage genes in the genomes of these *V. fischeri* cells (data not shown). To identify mutation(s) involved in resistance to ϕHNL01, we performed whole-genome re-sequencing on these ϕHNL01-resistant clones. A mutation shared among these resistant clones was a missense mutation encoding an E183V substitution in the glycosyltransferase gene VF_0175, which is part of the locus (VF_0157-80) responsible for production of exopolysaccharide (EPS) by strain ES114 (Bennett et al., 2020). Homologs of VF_0175 are predicted in many, but not all, *V. fischeri* genomes, suggesting that this gene may be under selection in different environmental contexts (Bongrand et al., 2020). Targeted deletion of VF_0175 or VF_0157-80, together with the provision of VF_0175 or VF_0175(E183V) *in trans,* confirmed that a functional copy of this gene in the context of a full-length EPS locus is necessary for ϕHNL01 infection (**Figure 3C**). We will subsequently refer to the VF_0175(E183V) “phage resistance” allele as VF_0175^PR^.

### *V. fischeri* EPS is associated with *ϕ* HNL01 susceptibility

To evaluate the extent to which the protein encoded by VF_0175 and the wild-type E183 residue contribute to EPS production, we used Alcian blue to stain EPS in liquid culture supernatants of *V. fischeri* ES114-derived strains. Targeted deletion of VF_0175 or VF_0157-80, along with providing VF_0175 or VF_0175^PR^ *in trans*, confirmed that, as was found for phage lysis, a functional copy of VF_0175 in the context of the full-length EPS locus is necessary for wild-type levels of EPS production (**Figure 3D**).

### Autonomous and squid-derived phage resistance contribute to symbiont colonization and maintenance

We next asked what impacts the VF_0175^PR^ allele had on competition with phage-sensitive *V. fischeri*. We co-cultured wild-type *V. fischeri* ES114 and its resistant VF_0175^PR^ derivative in the presence or absence of ϕHNL01, and found that in the presence of phage, wild-type ES114 was outcompeted by the VF_0175^PR^ strain, consistent with our observations of *V. fischeri* ES114 mutants in Figure 3 (**Figure 4A**). In addition, we observed that the VF_0175^PR^ strain monocolonized *E. scolopes* LOs as well as its wild-type parent *V. fischeri*, both in the presence and absence of phage in the ambient water (**Figure S2)**, suggesting that, rather than determining colonization ability, the natural variation in this locus is important for evading EPS-specific phages like ϕHNL01.

**Figure 4.**
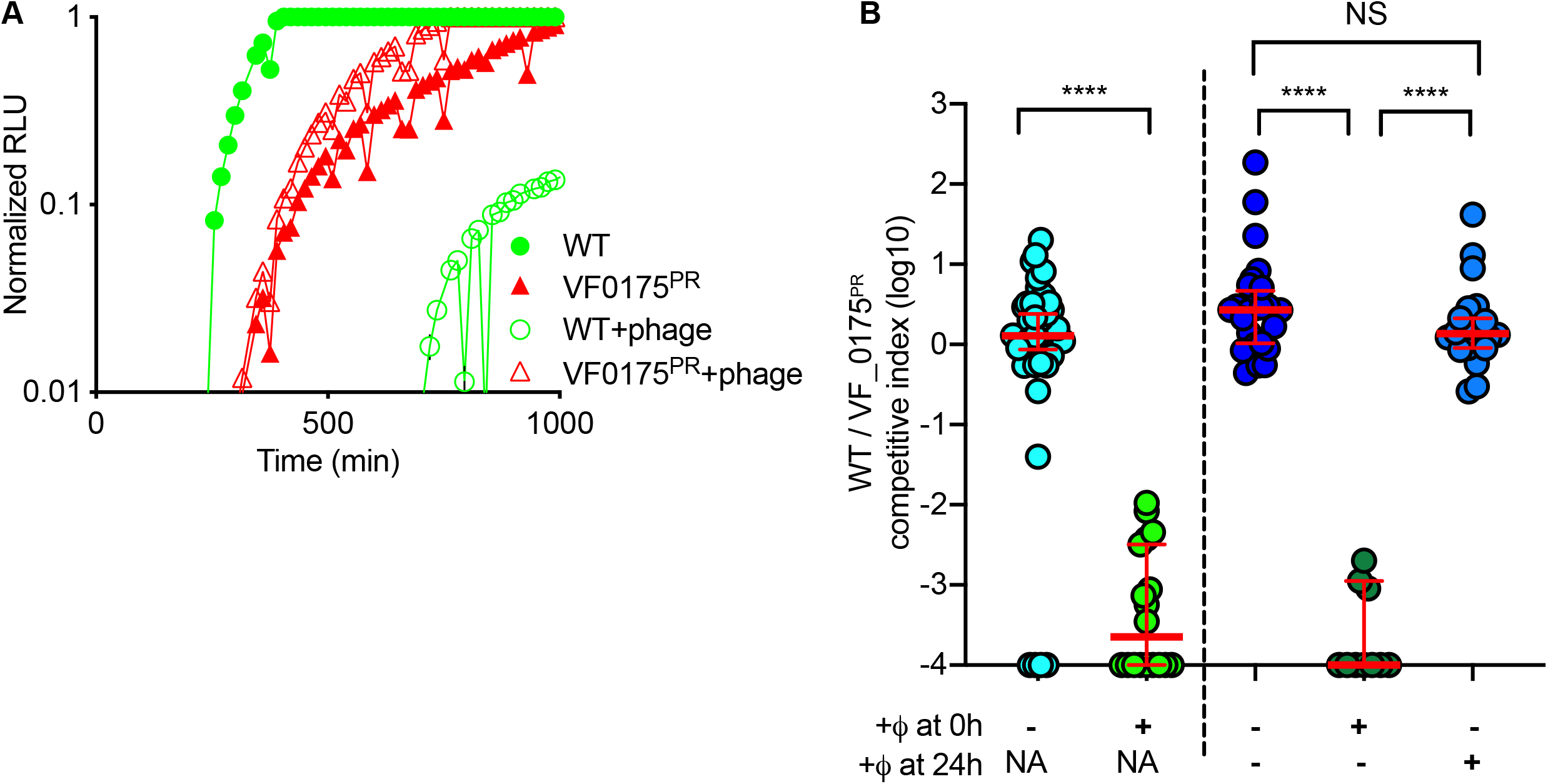
Innate phage resistance is beneficial to *V. fischeri* during planktonic lifestyle, but host provides protection after colonization. (A) Growth curves from cocultured differentially labeled wild-type *V. fischeri* ES114 and *V. fischeri* ES114 VF_0175^PR^ with or without phage exposure. WT was labeled with GFP and the VF_0175^PR^ mutant was labeled with RFP to distinguish them during co-culture:. Data presented are normalized RLU: (RLU – RLU_t=0_)/(maximum RLU range for that strain). Points represent mean +/−SEM. (B) Relative ratios of CFU from wild-type or strain VF_0175^PR^ from light organ homogenates of cocolonized squid. Phage were introduced at time = 0, time = 24 h, or neither, and homogenates were measured at t = 24 h (left two columns) or t = 48h (right three columns). n = 10-38 for each condition. Bars represent median +/− 95% CI.

To extend our previous observations that bacteria residing in the LO are protected from phage predation, we conducted co-colonization experiments using differentially labeled wild-type and resistant VF_0175^PR^ *V. fischeri*. Both strains colonized LOs similarly well in the absence of phage predation at both 24 h and 48 hours post colonization (**Figure 4B**; median wild-type/VF_0175^PR^ competition index = 1.28 at 24 h and = 2.68 at 48h). However, when squid were simultaneously exposed to bacteria and phage, as in **Figure 2B**, the wild-type parent was greatly outcompeted by the VF_0175^PR^ strain (**Figure 4B**; median wild-type/VF_0175^PR^ competition index = 2.24 x 10^−4^ at 24 h and <1 x 10^−4^ at 48h; p<0.0001 when compared with no phage at either time point, Mann-Whitney test) (**Figure 4B**). Finally, we conducted a sequential exposure experiment, where squid that were initially co-colonized with wild-type and VF_0175^PR^ were subsequently exposed to phage at 24 hours post inoculation. Though phage presence during the initiation of colonization led to the dominance of phage-resistant VF_0175^PR^ *V. fischeri* in the symbiosis, equal ratios of resistant and susceptible bacteria were present in LOs initially colonized without the presence of phage, regardless of later phage addition (**Figure 4B,** median wild-type: VF_0175^PR^ competitive index without phage = 2.68; median competitive index with phage after cocolonization = 1.37, p = 0.187 Mann-Whitney test).

## Discussion

Here we describe two synergistic means by which a mutualistic bacterium can be protected from phage predation: a cell-intrinsic, exopolysaccharide-based resistance factor and a host-derived shielding effect. Using the highly specific mutualism between *E. scolopes* and *V. fischeri*, we showed that these mechanisms protect the normal host-microbe association despite high symbiont clonality and a continuous connectivity between the symbiont culture and ambient seawater. We also showed that these types of protection against phage are important at different stages of colonization; bacterium-determined resistance is important during the transition from free living to the initial colonization, while symbiotic host-determined protection plays a greater role during the maintenance of an already established colonization. Together, these findings reveal several important principles of host-microbe-virus associations.

We were surprised that the newly isolated phage ϕHNL01 closely resembled ICP1, which is both highly prevalent in the stool of human cholera patients and may play a role in seasonality of cholera through selection against the *V. cholerae* O-antigen, the primary receptor for ICP1 (Seed et al., 2011, 2012). While we found that some membrane components are not required for ϕHNL01 predation (*e.g*., mutations in *waaL* and *eptA*, both of which are involved in lipooligosaccharide decoration, do not affect ES114 susceptibility to ϕHNL01; **Table S1**), we did identify other envelope-based infection resistance (*e.g*., mutations in the EPS gene cluster render ES114 resistant to ϕHNL01), suggesting some general shared aspects of phage infection between *V. cholerae* and *V. fischeri*. Because *V. fischeri* ES114 is the only ϕHNL01 host we identified, broader studies of *V. fischeri* phages around different areas of Hawaiʻi and other geographic regions where this species is endemic will reveal the ubiquity of related phages amongst different host populations, as well as the breadth of resistance mechanisms *V. fischeri* uses amongst its various symbiotic relationships (Bongrand et al., 2020).

Envelope and extracellular matrix modifications are major determinants of host-phage relationships at the individual and population levels (Díaz-Pascual et al., 2019; Dunsing et al., 2019; Hryckowian et al., 2020; Kim et al., 2019; Porter et al., 2020; Simmons et al., 2020), emphasizing the importance of surface factors in phage susceptibility and community dynamics. Resistance factors are important with respect to the survival of individual bacterial cells, but they likely become even more critical in areas with high densities of related and susceptible cells, where even inefficient infections could quickly sweep through a population. Scenarios like this include not only highly specific symbioses like the squid-*Vibrio* system, but it may also include spatially organized complex communities, such as biofilms and microcolonies.

Given the ubiquity of phages in the biosphere and their enrichment at mucosal surfaces (Barr et al., 2013, 2015), we asked whether associating with *E. scolopes* promoted infection of *V. fischeri* by phages, perhaps as a regulator of symbiont density or function. Instead, our work demonstrated that *E. scolopes* is a safe haven from viral infection, and colonization protected *V. fischeri* from phage predation. While this host-mediated protection may most directly benefit *V. fischeri*, it also is likely to benefit the squid by protecting its mutualist asset. This type of shielding may be especially important when a specific bacterial symbiont is not easily replaceable, either because of a colonization refractory period, niche competition, or ecological dynamics with the rest of the microbiota (Koch et al., 2014; Rao et al., 2021; Speare et al., 2018), as has been reported with the squid–*Vibrio* symbiosis. Similar observations have been made in gnotobiotic mice, where the distribution of gastrointestinal bacteria in phage-accessible and phage-inaccessible sites was hypothesized to be a driver of the long-term co-existence of phage and bacteria (Lourenço et al., 2020). Conversely, while symbiont protection in mutualism could benefit the host, it could similarly backfire during pathogenesis: surface-located bacterial pathogens would also be protected from phage, potentially even ameliorating the effectiveness of medical phage therapies that target these infections.

While the exact means by which the squid protects *V. fischeri* remains unclear, we speculate that several processes with broader implications may play a role. For one, the epithelium in the LO is proficient at uptake of small particles, which might allow non-specific consumption or transport of phage before they can infect *V. fischeri* in the crypts, as has been shown with other eukaryotic cells (Bichet et al., 2021; Cohen et al., 2020; Nguyen et al., 2017). *E. scolopes* generates flow fields through the beating of cilia on the exterior of the LO, preferentially promoting aggregation of *V. fischeri-*sized particles around the LO entry points, and it is also possible that these flow patterns may actively exclude phage-sized particles from the LO surface, mechanically protecting *V. fischeri* as it begins the process of colonizing the squid (Nawroth et al., 2017). These processes would hold broadly applicable implications for phage infections in other intimate symbioses, such as the mammalian gastrointestinal tract, where hosts might nonspecifically protect epithelium-adjacent symbionts.

This work also shows that in *V. fischeri*, mutation-derived resistance to phage readily appears under phage predation (**Figure 3A, B**). In four independent ϕHNL01-resistant clones, we identified mutations in VF_0175, which encodes a putative glycosyltransferase. While we did not identify any fitness costs imposed by the VF_0175^PR^ mutation during *in vitro* growth or colonization, it is possible that such naturally arising strains would normally be outcompeted by wild-type *V. fischeri* in the absence of phage, but random or specifically induced mutation to VF_0175 is strongly selected for during phage attacks. There are other examples of genetically acquired responses to phage infection that lead to bacterial resistance (e.g., CRISPR, mutation hot spots), and further work may uncover whether this high level of mutation is due to random mutation and selection or a more targeted genetic mechanism.

VF_0175 is necessary for plaque formation on *V. fischeri* ES114, but introducing *VF_0175* in the absence of the surrounding *VF_0157-0180* locus does not confer phage sensitivity, suggesting that carriage of a VF_0175 homolog is not sufficient for infection (**Figures 3C**). Interestingly, these mutant strains produce a lower level of EPS than wild-type ES114 (**Figure 3D**), suggesting that reduced EPS production itself may be selected for by ϕHNL01-like phages. Another EPS type produced by *V. fischeri*, the *syp* polysaccharide, is required during LO aggregation and colonization (Yip et al., 2006), demonstrating the delicate balance between selective factors that *V. fischeri* faces in symbiosis. Importantly, EPS production appears to be upregulated by genes expressed by *V. fischeri* in the LO (Bennett et al., 2020), where they would be at high density and most susceptible to phage infection, highlighting the potential importance of host protection at this critical point in the symbiosis.

Using a binary symbiotic system, we show that host- and bacteria-derived features can protect a host-microbe mutualism in the face of phage exposure. This work illuminates key approaches that members of a eukaryote’s microbiota can use to escape or avoid phage predation, providing a resilience mechanism for these communities. This work answers longstanding questions about the maintenance of highly specific symbiont populations, and serves as a foundation for investigating parameters that lead to protection in the LO. More broadly, this helps us to understand rules that modulate phage susceptibility in a specific host-associated niche, a key unanswered question necessary for harnessing phages as therapeutic agents and understanding the role of phages in microbiota dynamics.

## Methods

### Lead contact

Further information and requests for resources and reagents should be directed to and will be fulfilled by the Lead Contact, Andrew Hryckowian.

### Materials availability

ϕHNL01 and strains generated as part of this study are available upon request from the Lead Contact and from Edward Ruby, respectively.

### Data and code availability

During peer review, the .fastq files used to assemble the genomes of ϕHNL01, wild type *V. fischeri* ES114, and the 4 spontaneous ϕHNL01-resistant mutants of *V. fischeri* ES114 are available using the following Google Drive link. This link allows anonymous access to the files: https://drive.google.com/drive/folders/1TC00B6J8duD9oMcYAV3yP8HB649do7-E?usp=sharing.

Genomes are uploaded to NCBI and will be made freely available upon acceptance of the peer-reviewed manuscript.

### Animal husbandry

Adult *Euprymna scolopes* were collected from the waters of Maunalua Bay, Oʻahu, Hawai‘i, and maintained at Kewalo Marine Laboratory, where they would lay eggs as previously described (Montgomery and McFall-Ngai, 1993). Juvenile squid were collected <3 hours after hatching and placed in filter-sterilized ocean water (FSOW) and maintained on a 12:12-h light-dark photic cycle for the duration of experiments.

### Bacterial strains and culture conditions

The light organ symbiont *V. fischeri* strain ES114 (Boettcher and Ruby, 1990) was used for all experiments unless otherwise noted. Bacteria were grown in Lysogeny broth-salt medium (LBS) (10 g tryptone L^−1^, 5 g yeast extract L^−1^, 342 mM NaCl, and 20 mM Tris, pH 7.5) (Stabb et al., 2001) or seawater-tryptone medium (SWT) (70% ocean water, 32.5 mM glycerol, 5 g tryptone L^−1^, 3 g yeast extract L^−1^) (Boettcher and Ruby, 1990) at 28 °C, either on agar plates or in liquid culture shaking at 225 rpm. *Escherichia coli* WM3064 was grown in lysogeny broth (Bertani, 1951) supplemented with 60mM diaminopimelic acid at 37°C. Where appropriate, chloramphenicol (2.5 μg mL^−1^) or kanamycin (50 μg mL^−1^) was added to the media.

### Isolation of *Vibrio fischeri*-infecting bacteriophage ϕHNL01

Approximately 700 mL of coastal ocean water was collected each from Kewalo Basin and Kāneʻohe Bay on Oʻahu, Hawai‘i, USA, and was centrifuged at 5,500 x g for 10 minutes at room temperature to precipitate solids. The supernatants were then sequentially filtered through 0.45 μm and 0.22 μm pore polyvinylidene fluoride (PVDF) filters. The filtered water was concentrated 750-fold via 100 kDa PVDF size exclusion columns. The initial screening of plaques was performed using a soft agar overlay method in which 35 μL of filtered and concentrated seawater was combined with 200 μL of overnight *V. fischeri* ES114 LBS culture and with 4.5 mL molten SWT top agar (cooled to ~50°C, 3.5 g L^−1^ agar) and poured onto a warmed SWT agar plate (15 g L^−1^ agar).. Soft agar overlays were incubated aerobically at room temperature overnight.

Single isolated plaques were picked into 100 μL autoclaved Instant Ocean (IO; Spectrum Brands, Madison, WI, USA), and serial dilutions were prepared and spotted onto a solidified top agar overlay. Isolated plaques were picked after overnight growth. This procedure was repeated for a total of 3 times to purify ϕHNL01.

A high-titer stock of ϕHNL01 was generated by flooding a soft-agar overlay plate that yielded a ‘lacy’ pattern of bacterial growth (near confluent lysis). After overnight incubation of the plate, 5 mL of sterile IO were added to the plate to resuspend the phage. After at least 2 hours of incubation at room temperature, the lysate was extracted and filter sterilized through a 0.22 μm PVDF filter.

### Plaque assays

Isolated colonies from freshly streaked −80°C *V. fischeri* stocks were used to inoculate liquid LBS cultures. After ~16 h, 200 μL of culture were mixed with 4 mL molten SWT top agar and poured onto warmed SWT agar plates. 1 μL of serial 1:10 dilutions of phage lysate was spotted onto solidified top agar, and plates were incubated at 28°C for 3-5 h before counting plaques. Alternatively, 500 μL of bacteria (either from a stationary-phase culture of *V. fischeri* in SWT or pooled squid homogenates) were added to 4.5 mL of molten SWT top agar, then spread onto a warmed SWT plate and allowed to solidify. 5 μL of diluted phage (10^9^ PFU mL^−1^ unless otherwise noted) were spotted onto plates, and after drying, plates were incubated at 28 °C for 24 hours before checking for lysis. PFU counts and phage susceptibility assays were performed on SWT-agar plates, as plaques were noticeably smaller or non-observable on LBS plates.

### ϕHNL01 genome sequencing

DNA was extracted from a high-titer ϕHNL01 lysate and sequencing libraries were prepared using the Ultra II FS Kit (New England Biolabs, Ipswich, MA, USA). Libraries were quantified using a BioAnalyzer (Agilent, Santa Clara, CA, USA) and subsequently sequenced using 150-base single-end reads (Illumina MiSeq). Reads were imported into Geneious Prime (2021.1.1), quality-trimmed at an error rate of 0.001%. The Geneious assembler was used to assemble 30% of the surviving reads ≥150bp, with the options “Medium sensitivity / Fast” and “circularize contigs” selected. Coverage for the genome assembly reported by Geneious was 170 +- 21.

### ϕHNL01 genome annotation and comparative analysis with *Vibrio cholerae* phage ICP1

A 383-bp short direct terminal repeat was identified in ϕHNL01 by visualizing read pileups in Geneious Prime (2021.1.1), and the genome was arranged to place this repeat at the 5’ end of the genome. Protein-coding genes and tRNAs were predicted and annotated using DNA Master default parameters (http://cobamide2.pitt.edu), which incorporates Genemark (Besemer and Borodovsky, 2005), Glimmer (Delcher et al., 1999), and tRNAscan-SE (Lowe and Eddy, 1997). Phage genomes were annotated and compared on the basis of shared gene phamily (pham) membership within Phamerator using default parameters (Cresawn et al., 2011). Phams are groups of related protein-encoding genes where pham membership is built and expanded when a candidate protein shares ≥32.5% identity or a BLASTp e-value <1e^−50^ with one or more existing members of the pham. Genome maps of ϕHNL01 and ICP1 were visualized in Phamerator using default parameters. Amino acid sequences for ϕHNL01 and ICP1 were concatenated and visualized as a dotplot using word size 5 in Geneious Prime (2021.1.1)

### Plasmid and mutant construction

Primers used to create the *V. fischeri* gene deletion and expression plasmids used in this study are listed in **Table S3** and **Table S1**, respectively. In-frame deletion of *VF_0175* from the *V. fischeri* genome was performed as previously described (Bennett et al., 2020; Saltikov and Newman, 2003). Briefly, ~1kb fragments surrounding *VF_0175* were fused via an internal *Eco*RI site and inserted via *Bam*HI and *Sac*I sites into pSMV3 (Coursolle and Gralnick, 2010), which has kanamycin resistance and *sacB* cassettes. Plasmids were introduced to *V. fischeri* ES114 through conjugation with *Escherichia coli* WM3064, and counter-selection to remove pSMV3 and VF_0175 was performed on low-salt, high-sucrose LB plates (Bennett et al., 2020) at room temperature. The wild-type *VF_0175* complementation plasmid was constructed by cloning *VF_0175* from the *V. fischeri* ES114 genome, and inserting it into the expression plasmid pVSV105 (Dunn et al., 2006) via *Sph*I and *Kpn*I sites. The VF_0175^PR^ allele was ordered as a gblock from Integrated DNA Technologies (Coralville, IA, USA), and inserted into pVSV105 via *Sph*I and *Kpn*I sites.

### Alcian blue detection of extracellular polysaccharides

EPS was detected as previously described (Bennett et al., 2020). Briefly, overnight bacterial cultures grown in minimal salts medium supplemented with 6.5 mM *N*-acetylneuraminic acid and 0.05% (wt/vol) casamino acids were centrifuged for 15 min at 12,000 x g and 4 °C. 250 μL culture supernatant was incubated with 1 mL Alcian blue reagent on a rocker for one hour at room temperature, then centrifuged 10 min at 10,000 rpm and 4 °C. Pellets were washed with 1 mL 100% ethanol and centrifuged 10 min at 10,000 x g and 4 °C. Pellets were solubilized in 200 μL SDS-acetate and the absorbance read at OD_620_.

### Identification of phage-resistance genes in *Vibrio fischeri*

Four colonies from were picked from the center of ϕHNL01 phage spots containing at least 10^5^ plaque forming units (PFU). Colonies were streaked for isolation over two more passages on LBS plates. Isolates were grown overnight in LBS medium and were used to make glycerol stocks and confirm phage insensitivity. An overnight LBS culture was used for DNA extraction using the Qiagen Blood and Tissue Kit (Qiagen, Germantown, MD, USA). Sequencing libraries were prepared from genomic DNA from each ϕHNL01-resistant isolate, as well as the parental *V. fischeri* ES114 strain, using the Ultra II FS Kit (New England Biolabs). Libraries were quantified using a BioAnalyzer (Agilent) and subsequently sequenced using 151-base single-end reads (Illumina MiSeq). Differences between wild-type and resistant *V. fischeri* ES114 was identified through genomic comparison using CLC Genomics Workbench (Qiagen).

### *Vibrio fischeri* colonization and phage exposure

Juvenile squid were incubated with ~10^4^–10^5^ colony-forming units (CFU) mL^−1^ of wild-type *V. fischeri* ES114 and/or ϕHNL01-resistant ES114 VF_0175^PR^ (as noted) for 24 hours. For cocolonizations, ~1:1 inocula of each strain were used, and competitive index was measured relative to strain ratio in the inoculum (competitive index = (wildtype:mutant ratio in LO)/(wild-type: mutant ratio in inoculum). When appropriate, animals were transferred to fresh FSOW after 24 hours. Animals were frozen at −80°C in FSOW immediately before dark (*i.e*., at maximal colonization), then thawed and homogenized, and serial dilutions were plated on LBS-agar plates to determine *V. fischeri* CFU counts. GFP- and RFP-positive CFU were counted using a fluorescence dissecting microscope.

When appropriate, phage was added to the water at ~10^7^ PFU mL^−1^.

### In vitro *phage exposures*

ϕHNL01-sensitive and resistant *V. fischeri* ES114 strains received pVSV102 (GFP, Kan^r^), pVSV208 (RFP, Cam^r^), or pVSV104-H (Kan^r^) (Dunn et al., 2006) via conjugation with donor strain *Escherichia coli* WM3064 (Lynch et al., 2019). Cultures of wild-type and/or ϕHNL01-resistant *V. fischeri* ES114 were streaked onto LBS plates with the appropriate antibiotic and then cultured in LBS overnight. Cells were then diluted in SWT as mentioned in the experiment. 100μL of diluted cells were placed in wells of a 96-well flat bottom transparent plate with or without ϕHNL01 at the noted dilutions of a ~10^11^ PFU mL^−1^ stock, and were incubated at 28 °C with periodic shaking for 24-48 hours. A Tecan GenIOS plate reader (Tecan, Baldwin Park, CA, USA) was used to measure GFP and RFP fluorescence and/or overall cell density (OD_600_).

### PCR detection of phage genes

Taq DNA polymerase was used perform to PCR against ϕHNL01-specific genes (*gp47*, *gp61*, *gp96*, *gp147;* see **Table S3**) on glycerol stocks of ϕHNL01-resistant *V. fischeri* isolates to assess their potential lysogen status (see **Table S3** for primer sequences). Cycling conditions were: 95 °C for 5 min, 35 cycles of 95 °C for 30 sec→ 55 °C for 30 sec→ 68 °C for 1 min, then 68 °C for 5 min followed by a 4°C hold. PCR was also run directly on phage samples and on wild-type ES114 as controls. Products were run on a 1% agarose gel to determine PCR product presence/absence, and results were confirmed by genome sequencing.

### Electron microscopy

Serial dilutions of phages were spotted onto *V. fischeri* ES114 + SWT top-agar lawns as described above and allowed to form plaques overnight. Dilutions with the most easily distinguishable plaques (*i.e*., the lowest dilution that still produced plaques) were briefly covered with 10 μL of FSOW to resuspend phage, then removed to a clean PCR tube. A 4 μL portion of the suspension was spotted onto a charged formvar-coated copper grid, negatively stained with uranyl acetate, and imaged with a Hitachi HT7700 transmission electron microscope at 100kV at the University of Hawaiʻi MICRO Facility.

### Quantification and statistical analysis

Statistical analysis was performed using Graphpad Prism 9.1.0. Details of specific analyses, including statistical tests used, are found in applicable figure legends. All experiments were performed in at least triplicate. * = p<0.05, ** = p<0.005, *** = p<0.0005, **** = p<0.0001.

## Acknowledgements

We thank Randall Scarborough for logistical assistance in collecting seawater samples; Daniel Russell and Rebecca Garlena for sequencing phage and bacterial genomes; Tina Carvalho for assistance with transmission electron microscopy. This work was funded by NIH grant F32 GM119238 and a Ford Foundation Postdoctoral Fellowship to J.B.L., a National Science Foundation Graduate Research Fellowship DGE-114747 to B.D.M., NIH R01 GM135254 and R01 AI050661 to E.G.R., and startup funding from the University of Wisconsin-Madison to A.J.H. Microscopy was performed at the University of Hawaiʻi’s MICRO facility, supported under COBRE grant P20 GM125508.

## Author contributions

J.B.L., B.D.B., B.D.M., and A.J.H., performed experiments/computational analyses and analyzed the data. J.B.L., B.D.B., B.D.M., and A.J.H. prepared the display items. E.G.R. provided key insights, tools, and reagents. J.B.L., B.D.B., and A.J.H. wrote the paper. All authors edited the manuscript prior to submission.

---

**Figure S1. Genome maps of ϕHNL01 and ICP1.** Genes are represented as colored boxes and conserved domains are inlaid yellow boxes within genes. If a gene has a conserved domain, it is annotated in black text. Pairwise nucleotide identity is represented as shading between genomes. The color of this shading represents the degree of sequence similarity with violet being the most similar (BLASTN score=0), progressing through the color spectrum from indigo, blue, green, yellow, orange, to red, which is the least similar (BLASTN score = 10^−4^). Regions with no shading indicate no similarity with a BLASTN score greater than 10^−4^. Related to Figure 1.

**Figure S2. Phage exposure and phage resistance does not affect overall colonization levels of wild-type *V. fischeri* ES114 or *V. fischeri* ES114 VF_0175^PR^.** CFU LO^−1^ from squid monocolonized with wild-type *V. fischeri* ES114 or phage resistant *V. fischeri* ES114 VF_0175^PR^ with or without phage. Related to Figure 4.

**Table S1. Bacterial strains and plasmids used in this study.** Related to Figures 1, 2, 3, and 4.

**Table S2. Conserved domains identified between ϕHNL01 and ICP1.** Related to Figure 1.

**Table S3. Primers used in this study.** Related to Figures 3 and 4.

