## Supplementary figures and images for "Independent host- and bacterium-based determinants protect a model symbiosis from phage predation"

### Supplemental Figure 1

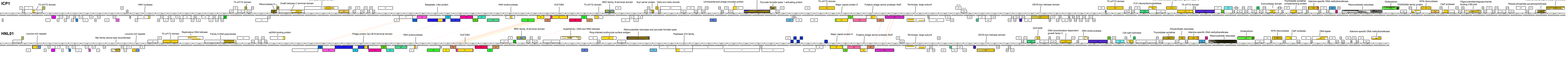

### Supplemental Figure 2

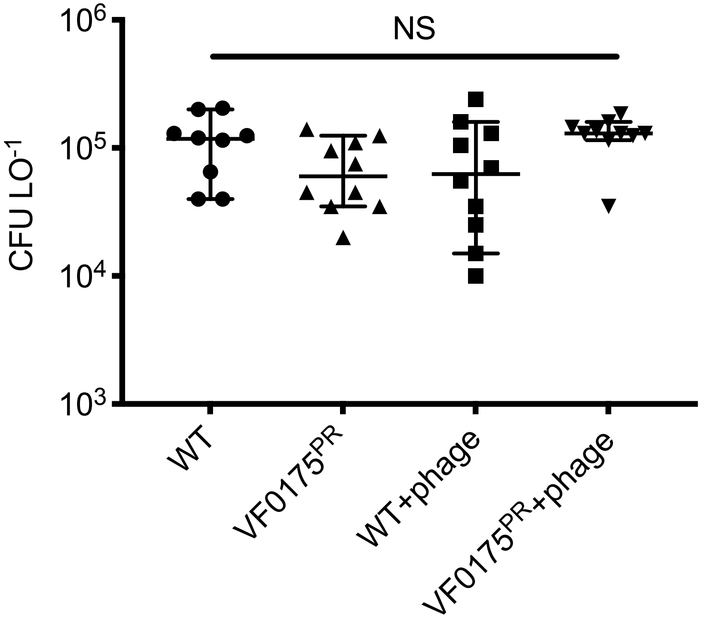
